# A viral guide RNA delivery system for CRISPR-based transcriptional activation and heritable targeted DNA demethylation in *Arabidopsis thaliana*

**DOI:** 10.1101/2020.07.09.194977

**Authors:** Basudev Ghoshal, Brandon Vong, Colette L. Picard, Feng Suhua, Janet May Tam, Steven E. Jacobsen

## Abstract

Plant RNA viruses are used as delivery vectors for their high level of accumulation and efficient spread during virus multiplication and movement. Utilizing this concept, several viral-based guide RNA delivery platforms for CRISPR-Cas9 genome editing have been developed. The CRISPR-Cas9 system has also been adapted for epigenome editing. While systems have been developed for CRISPR-Cas9 based gene activation or site-specific DNA demethylation, viral delivery of guide RNAs remains to be developed for these purposes. To address this gap we have developed a tobacco rattle virus (TRV)-based single guide RNA delivery system for epigenome editing in *Arabidopsis thaliana*. Because tRNA-like sequences have been shown to facilitate the cell-to-cell movement of RNAs in plants, we used the tRNA-guide RNA expression system to express guide RNAs from the viral genome to promote heritable epigenome editing. We demonstrate that the tRNA-gRNA system with TRV can be used for both transcriptional activation and targeted DNA demethylation in Arabidopsis. We achieved up to ~8% heritability of the induced demethylation phenotype in the progeny of virus inoculated plants. We did not detect the virus in the next generation, indicating effective clearance of the virus from plant tissues. Thus, TRV delivery, combined with a specific tRNA-gRNA architecture, provides for fast and effective epigenome editing.

**Author summary:** The discovery of CRISPR-CAS9 and its non-catalytic variants have provided enormous capacity for crop improvement and basic research by modifying the genome and the epigenome. The standard methods for delivering genome and epigenome editing reagents to plants consist of generating stable transgenic lines through tissue culture processes, which have several drawbacks including the need for plant regeneration and crossing. To overcome some of these challenges, plant virus-based platforms are being developed for genome editing. Although viruses have a limited cargo capacity, limiting the use of viruses to encode entire editing systems, guide RNAs have been successfully delivered to transgenic CAS9 expressing plants for genome editing. However, the use of viruses for CRISPR-based epigenome editing and transcriptional activation have not yet been explored. In this study we show that viral delivery of guide RNAs using a modified tobacco rattle virus can be used for transcriptional activation and heritable epigenome editing. This study advances the use of plant RNA viruses as delivery agents for epigenome editing.

## Introduction

Tools that enable targeted regulation of gene expression provide powerful systems for studying and manipulating diverse cellular processes [1]. Targeted gene regulation at the transcriptional level can be achieved by recruiting transcriptional activators, repressors, or epigenetic modifiers to a particular genomic locus using a programmable DNA-binding module, such as the catalytically inactive version of the CRISPR-CAS9 system [1–3]. The CRISPR-CAS9 system is comprised of the CAS9 endonuclease and single guide RNA (gRNA) [4, 5]. The modularity and simplicity of this system to target many sites of the genome have made it a popular tool for targeted genome editing. Catalytically inactive CAS9 (dCas9) can act as a DNA binding module, enabling the targeted recruitment of any effector protein attached to dCas9. Such CRISPR-dCAS9 based tools have been used to regulate gene expression and/or modify the epigenome across multiple species [3, 6–9].

The CRISPR-based SunTag system is a robust system for transcriptional gene regulation that is capable of recruiting multiple copies of an effector protein [6, 9, 10]. In this system, dCAS9 is fused to a chain of peptide epitopes, while effector proteins are separately fused to single-chain variable fragments (scFV). The scFVs bind to the epitopes attached to dCas9, allowing dCAS9 to recruit multiple copies of the effector proteins simultaneously to any sites targeted by the gRNA [9]. Using this system with the transcriptional activator VP64 led to robust upregulation of targeted genes and corresponding phenotypic changes in *Arabidopsis thaliana* [6]. Similarly, the SunTag system has been used to add or remove DNA methylation through site-specific recruitment of epigenetic modifiers, including the DNA methyltransferase DRM2 (DOMAINS REARRANGED METHYL TRANSFERASE 2) and the human demethylase TET1 (TEN ELEVEN TRANSLOCATION 1) [6, 8, 10]. SunTag-mediated epigenomic modifications can cause changes in both gene expression and the associated phenotype by removing DNA methylation from the *FWA* (*FLOWERING WAGENINGEN*) promoter. Researchers were able to express this gene constitutively in vegetative cells, thus delaying flowering of Arabidopsis plants [11, 12]. These targeted epigenetic modifications, and subsequent transcriptional changes, were found to be heritable and were maintained in the absence of the SunTag construct across several generations [6, 10]. CRISPR-based epigenome editing and transcriptional activation in plants thus has great potential for manipulating plant traits.

Plant viruses have been used to deliver components of the CRISPR-CAS9 genome editing platform owing to their high levels of accumulation and efficient spread to most plant cells [13, 14]. However, both plant RNA and DNA viruses have limitations, including limited cargo capacity and restricted entry into the plant meristem or the germline. The CRISPR-CAS9 system itself consists of a large DNA construct (~ 3 – 4 kb), which is larger than the cargo capacity of most viruses. The limitation in cargo capacity means that viral delivery of the entire editing system remains challenging in plants. Therefore, the viral-based delivery systems currently available for CRISPR-CAS9 genome editing systems are mainly focused on the delivery of guide RNA component of the system. Viral delivery has advantages over commonly used procedures of generating transgenic lines expressing Cas9 and guide RNAs. For example, a lower concentration of guide RNA is a limitation for efficient genome editing, which can be overcome by using viruses [15]. Secondly, viral delivery of guide RNAs to transgenic CAS9 plants may provide the advantage of bypassing the repeated need to generate transgenic plants expressing different guide RNAs each time a new gene is targeted. The generation of stable transgenic lines for many crop species is a time-consuming step involving tissue culture. Several studies have demonstrated the delivery of guide RNAs using different viruses for genome editing. Geminiviruses, which are single stranded DNA viruses, have been used to deliver both CAS9 and single guide RNAs [16, 17]. However, for the delivery of CAS9, a movement defective geminivirus was used [16]. Similarly, plant RNA viruses, including the Tobacco rattle virus (TRV), Tobacco mosaic virus, Barley Stripe virus, have been used to deliver gRNAs to plants [15, 18–20].

Despite the advantages of viral delivery of guide RNAs and the expansion of CRISPR based gene regulatory systems, currently, viral guide RNA delivery tools have thus far not been used for targeted epigenome editing. Here we extend the use of plant RNA viruses as vectors to deliver guide RNA for gene expression control and epigenome editing. We developed a viral vector system that can infect and multiply within plant cells and efficiently spread to germ cells to promote heritable epigenome editing. We utilized TRV, which infects a wide range of plant species including, Arabidopsis. TRV can transiently infect the shoot apical meristem [21], which makes it well suited as a viral vector for heritable genome [19, 20] or epigenome editing. Furthermore, to facilitate heritable epigenome editing, we incorporated tRNA-like mobile signal sequences in our vector. Certain plant RNA sequences, such as specific FT (*Flowering locus T*) and tRNA sequences, have been shown to act as mobile signals that facilitate the movement of endogenous non-mobile RNAs or viral RNAs within plants and into the meristem or the flowers [22, 23]. Very recently, Ellison et al. demonstrated for genome editing purposes in *Nicotiana benthamiana*, that the use of mobile signal sequences (FT and tRNA) in combination with TRV and guide RNAs enhanced the efficiency of heritability of edits [20]. Here we show that this “tRNA-guide RNA architecture”, in combination with TRV-mediated delivery, can be an effective strategy for epigenome editing using the SunTag system. Our results show that this viral delivery system can induce efficient heritable epigenome editing in Arabidopsis. The induced phenotype and gene expression changes are stable in subsequent generations and persist after the virus itself has been cleared.

## Results and discussion

### Transcriptional activation of *FWA* by TRV-delivered guide RNAs utilizing a tandemly arrayed “tRNA-gRNA architecture”

We developed a “tRNA-gRNA architecture” based TRV guide RNA delivery system for epigenome editing and gene activation. CRISPR-based transcriptional activation assays allow robust and precise assessment of mRNAs, the product of transcription. We therefore performed an initial set of experiments to test whether this TRV based guide RNA delivery system could be used for transcriptional activation. To this end, we utilized transgenic *Arabidopsis thaliana* plants expressing the SunTag system, with VP64 as the effector protein [6], but lacking the gRNA component of the system (S1a Fig). The *FWA* gene is a good target for transcriptional activation studies as it is normally transcriptionally repressed in vegetative cells [6]. We cloned a single gRNA (guide RNA4) targeting the *FWA* promoter flanked by two tRNAs into RNA2 of the bipartite TRV RNA genome for the proper expression of the guide RNA (S1b Fig). In this context, the subgenomic RNA promoter of the Pea early Browning virus (sgPeBV) drives tRNA-gRNA expression (S1b Fig). Two lower leaves of 12-14 day old *Arabidopsis thaliana* were infected with TRV containing the guide RNA 4 or with control TRV with no guides, propagated first in *N. benthamiana* plants. For simplifying naming of the constructs, hereafter, TRV with no guide RNA sequences are referred to as “-gRNA”, and TRV with guide RNA sequence are referred to as “+gRNA”, unless otherwise noted. TRV inoculated rosette leaves from 3 individual plants were collected at 3 days post-inoculation (dpi) and tested for *FWA* activation. We observed *FWA* activation ranging from thirty to seventy-eight fold in the 3 dpi inoculated leaves relative to no-gRNA controls (Fig 1a, (left panel) : Sample 1a, 1b, 1c) suggesting that guide RNAs delivered using this system are functional and able to activate *FWA* in inoculated leaves.

**Fig 1.**
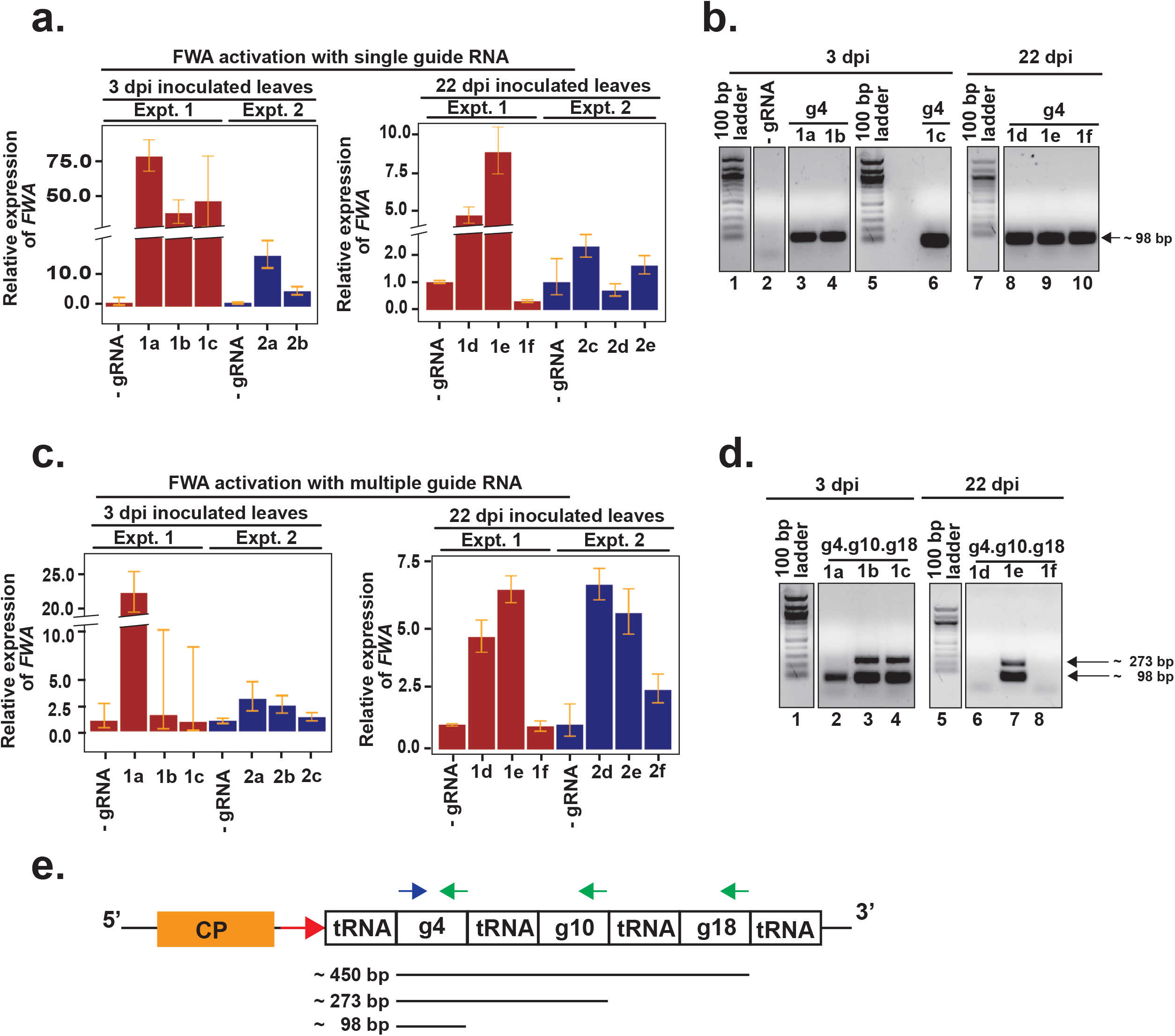
TRV-mediated guide RNA delivery in SunTag-VP64 plants causes transcriptional activation of *FWA*. Reverse transcriptase-qPCR was performed on total RNA extracted from leaves of individual plants at different time points post TRV inoculation to examine the accumulation of *FWA* mRNAs by single guide RNAs (a) and multiple guide RNAs (c). Fold expression change is relative to control plants inoculated with TRV with no guide RNA. Left panel shows *FWA* expression in samples collected from 3 dpi inoculated leaves and right panel shows samples from 22 dpi upper uninoculated cauline leaves in (a) and (c). Experiment 1 (Expt.1) and experiment 2 (Expt.2) indicate two independent biological experiments, each using multiple plants. Individual samples are denoted as “1a”, “2a” and so on. The prefix 1 or 2 denotes the experiment, the alphabets “a”, “b” denotes individual plants. Error bars indicate standard deviations (n=2 technical replicates for experiment 1, and n=3 technical replicates for experiment 2). To simplify labelling, TRV with no guide RNA is denoted as - gRNA, TRV with guide 4 as g4, and TRV with multiple guides as g4.g10.g18. Agarose gel electrophoresis of reverse transcriptase PCR products amplified to detect guide RNA specific sequences in g4 (b) and g4.g10.g18 (d) inoculated plants from experiment 1.100 bp band corresponds to guide 4 sequence (98bp), while 273 bp likely corresponds to unprocessed guide 4 sequence containing RNAs. Numbers at the bottom of the gels indicate lanes with samples or DNA ladder. Samples run on the lane numbers that are grouped here in parentheses (1-4), (5-6), (7-10) in (b) and (1-4), (5-8) in (d) were ran on the same gel. In addition, lane (5-6) in (b) and (1-4) in (d) were run on the same gel and so have a common marker. (e) Schematic representation of primer binding sites from (b) and (d) in the modified viral RNA2 containing three separate guide RNAs. Blue arrow indicates the forward primer, green indicates reverse primers. Red arrow downstream of Coat protein (CP) denotes the sgPeBV promoter. Expected sizes of PCR products are shown at the bottom.

To examine whether, TRV harboring guide RNAs were able to move systemically and cause transcriptional activation in the upper systemically infected leaves we collected uninoculated, upper cauline leaves at 22 dpi and analyzed them for *FWA* activation and the presence of viral RNA. An up to nine-fold increase in *FWA* expression was observed in plants infected with +gRNAs relative to the control plants by reverse transcriptase qPCR (Fig 1a, (right panel) : Sample 1d, 1e, 1f). To test for the presence of viral RNA in 3 dpi and 22 dpi leaves, we used a reverse transcriptase-PCR followed by agarose gel electrophoresis assay using guide 4 sequence-specific primers (Fig 1b). A 98 bp band corresponding to the guide 4 was detected in both 3 dpi inoculated and 22 dpi upper uninoculated leaves (Fig 1b), indicating virus accumulation and spread. Activation of *FWA* in the upper un-inoculated leaves demonstrated that TRV-containing guide RNAs can systemically spread within plants and transcriptionally activate *FWA*. To check for the reproducibility of the system, we repeated the experiment independently and analyzed TRV inoculated plants for *FWA* activation. *FWA* activation was again observed in the TRV inoculated plants (Fig 1a). However, the maximum level of activation observed was up to 16 fold in 3 dpi inoculated plants (Fig 1a, (left panel) : Sample 2a, 2b) and 2.3 fold in the 22 dpi upper uninoculated leaves (Fig 1a, (right panel) : Sample 2c, 2d, 2e) indicating an overall lower level of activation relative to the first experiment. This variability in the level of *FWA* activation between the two different experiments could be due to several factors, including improper processing of the guide RNAs, the concentration of initial viral inoculum, and variability in the spread of the virus relative to plant health.

The tRNA-gRNA expression system provides the ability to express multiple guides from the same transcript. It has been shown that utilizing several guides for transcriptional activation may have some synergistic effects [3, 6, 24]. Therefore we also tested whether plants inoculated with TRV containing multiple guide RNAs can activate *FWA* expression in the inoculated and systemically infected leaves. We cloned in three guide RNAs (guide 4, guide 10 and guide 18), each flanked by tRNAs in the RNA2 (S1b Fig) that bind to different regions of *FWA* relative to the TSS (S1c Fig). Leaf samples were collected from 3 dpi inoculated and 22 dpi upper uninoculated leaves of individual TRV inoculated plants for analyzing the presence of viral RNA and *FWA* activation. At 3 dpi, one plant out of the three inoculated showed activation (22 fold change) (Fig 1c, (left panel) : Sample 1a). At 22 dpi, we observed a maximum of 6.5 fold activation (Fig 1c, (left panel) : Sample 1e). We tested for the presence of the viral RNA in these plants, in addition to the 98 bp guide 4 sequence-specific band, we detected a larger PCR product (~ 300 bp) (Fig 1d). These are likely PCR products amplified from unprocessed viral RNAs where the reverse primer can bind to the multiple guide RNA scaffolds leading to the amplification of multiple bands (Fig 1e). We repeated the inoculation in an independent experiment using multiple guides, this time observed a 2.5 to 3 fold activation at 3 dpi and 2.4 to 6.7 fold at 22 dpi uninoculated leaves (Fig 1c, (left panel) : Sample 2a, 2b, 2c, (right panel) : Sample 2d, 2e, 2f) indicating activation but to variable levels. Contrary to the general observation that the use of multiple guides shows synergism and enhances activation we did not observe any such effects on *FWA* activation in our study. This could be due to uneven processing of guide RNAs from a longer tRNA-guide RNA transcript in comparison to the smaller single guide 4 transcript. Nonetheless, our data demonstrate that the tRNA-gRNA system can be combined with TRV delivery to express functional guide RNAs that can activate an Arabidopsis gene.

### Guide RNAs delivered by TRV can demethylate and activate the *FWA* promoter

We next tested whether our system could efficiently induce targeted epigenome editing in Arabidopsis. The *FWA* promoter is heavily methylated and silent in wild type plants. Loss of DNA methylation at the promoter results in *FWA* expression, leading to a late-flowering phenotype [11, 12]. We, therefore, tested whether our gRNA delivery system could be used to trigger the demethylation of *FWA*. We obtained transgenic plants expressing the human DNA demethylase, TET1, with the SunTag system, but lacking the gRNA component of the system [6]. We inoculated the SunTag-TET1 plants (the V1 generation) with the same TRV constructs used with SunTag-VP64 (S1b Fig). To bypass the step of producing viral inocula in *Nicotiana benthamiana*, we directly agro-inoculated transgenic SunTag-TET1 plants with cDNA infectious clones of TRV [25]. Agro-inoculated leaves were collected at 3 and 6 dpi, and upper non-agro-inoculated cauline leaves were collected at 22 dpi.

We first analyzed DNA methylation levels at the *FWA* promoter in inoculated plants using a restriction-enzyme based DNA methylation assay (McrBC-qPCR) (S2a Fig). We selected the two plants (V1-2, V1-9) with the highest level of loss of methylation together with matched -gRNA controls (V1-1, V1-5), for further analysis using a higher resolution technique, bisulfite PCR sequencing, in which *FWA* regions were amplified from bisulfite treated genomic DNA, and the amplicons subjected to high throughput Illumina sequencing [6]. We focused on CG methylation because it is known to be the key type of methylation that correlates with *FWA* silencing [26] (Fig 2a). We observed only a moderate loss of CG DNA methylation at three regions near the *FWA* TSS (Fig 2a). We hypothesized that demethylation might be occurring in only a subset of cells in the sample, such that average methylation levels are not strongly affected. To test this, we identified reads that were fully unmethylated in either the CG or CHH context. The CHG context was not considered due to the scarcity of CHG sites in the regions tested. We calculated the fraction of fully unmethylated reads in each +gRNA sample, normalized to the same value in the matched -gRNA control. Both lines (V1-2, V1-9) showed a strong enrichment of fully CG-unmethylated reads compared to the controls, indicating DNA demethylation of CG sites in this region (Fig 2b). Interestingly, the same analysis showed no evidence of CHH methylation loss in these lines, consistent with TET1’s known preference for demethylating CG sites in Arabidopsis [27] (S2b and S2c Figs). These results are consistent with a recent study that found that overexpression of TET1 in Arabidopsis had severe effects on CG methylation, but not on CHG and CHH methylation [27].

**Fig 2.**
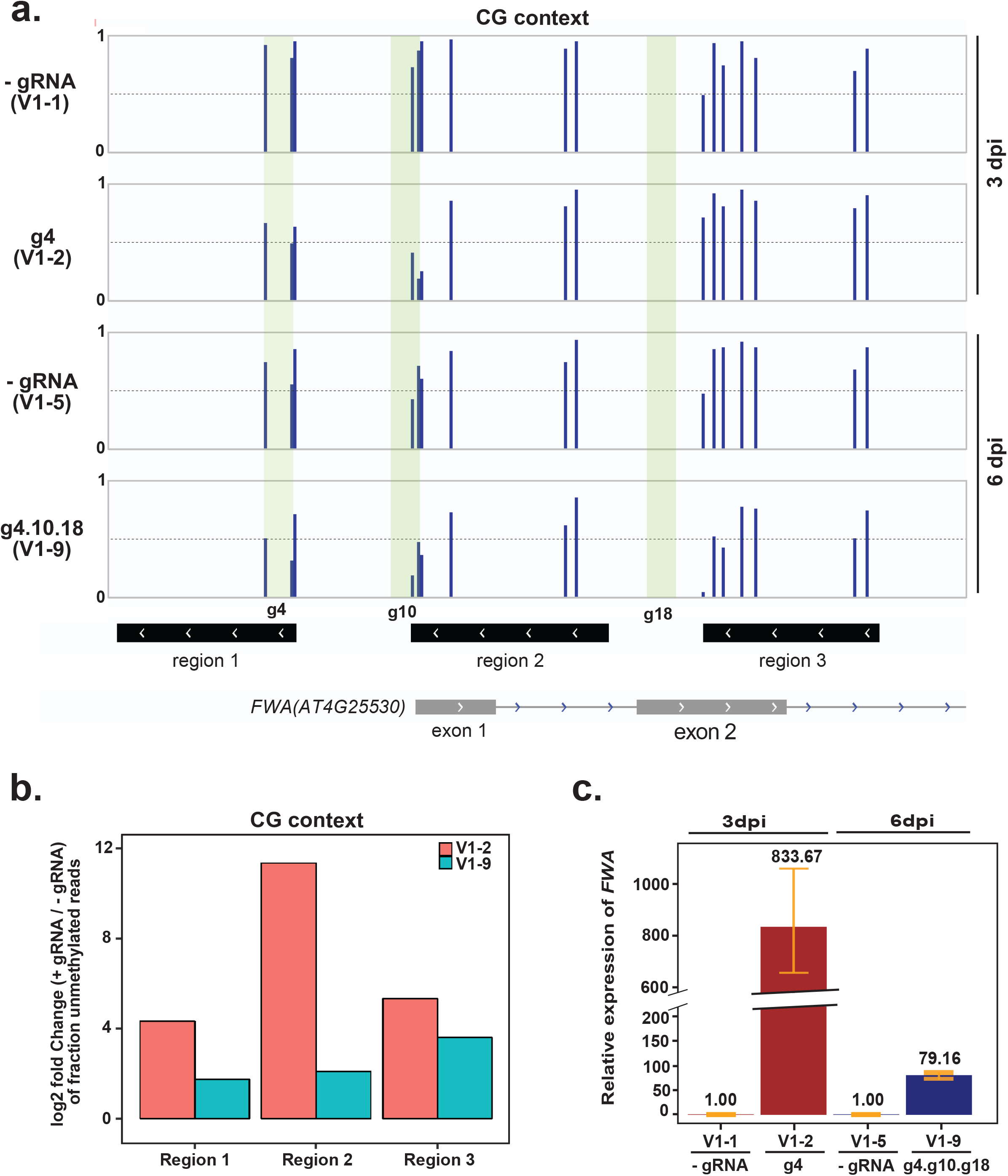
TRV-mediated guide RNA delivery in SunTag-TET1 plants triggers DNA demethylation and transcriptional activation of *FWA*. (a) Bisulfite PCR sequencing of two +gRNA lines (V1-2, V1-9) and their matched -gRNA controls (V1-1, V1-5) over three regions of the *FWA* promoter region: region 1 (Chr4:13038143 to 13038272), region 2 (Chr4:13038356 to 13038499) and region 3 (Chr4:13038568 to 13038695), using genomic DNA extracted from rosette leaves of TRV inoculated plants collected at 3 and 6 dpi. Only CG sites are shown. An average of 30,000 to 50,000 reads were obtained for each of the three regions. Green stripes denote guide4, guide 10 and guide 18 binding site. (b) Data shown in (a) was displayed as a bar graph of the log_2_ ratio of the fraction of fully CG-demethylated reads in +gRNA plants relative to their matched -gRNA controls. (c) *FWA* expression in +gRNA lines compared to control plants by reverse transcriptase-qPCR. Each +gRNA and its matched control were normalized so that expression in the control was equal to 1. Error bars indicate standard deviations (n=3 technical replicates). All comparisons were made with their respective matched controls. For example, 3 dpi sample + gRNA (V1-2) line was compared to -gRNA line (V1-1) of the same time point.

We next determined whether the loss of CG methylation observed in these plants led to an upregulation of *FWA* expression by reverse transcriptase-qPCR. Both plants (V1-2 and V1-9) showed a dramatic increase in the level of *FWA* expression, an 833- and 79-fold upregulation, respectively (Fig 2c). However, neither of these two plants exhibited a late-flowering phenotype, likely owing to the fact that TRV inoculation was performed on 12-14 day-old seedlings, which is after the flowering time decision has already been made [28].

### Guide RNAs delivered by TRV can trigger epigenetic changes in SunTag-Tet1 plants that are transmitted to progeny

To test whether TRV encoded gRNAs can affect the germline of V1 plants and cause heritable DNA demethylation and expression changes at *FWA*, we analyzed progeny of selfed V1 plants (the V2 generation), as well as subsequent generations, and adopted the naming system shown in S3a Fig. We selected approximately 50 V2 plants derived from each of the two V1 lines (V1-2, V1-9), and monitored the plants for a late-flowering phenotype. Several progenies of controls (V1-1, V1-5) inoculated with -gRNA were similarly selected and monitored. Once plants had flowered, we counted the total number of leaves on each plant, which is an indicator of flowering time. The V2 progeny of -gRNA plants had ten to sixteen leaves at the time of flowering, within the typical range of wild type, while a number of progeny of +gRNA inoculated plants had more than sixteen leaves, indicating delayed flowering time (S1a Table). Seven of these plants, or 6.6%, met our strict criteria for a late-flowering phenotype (22 leaves), while a number of other V2 plants showed an intermediate delayed flowering (Fig 3a and S1a Table).

**Fig 3.**
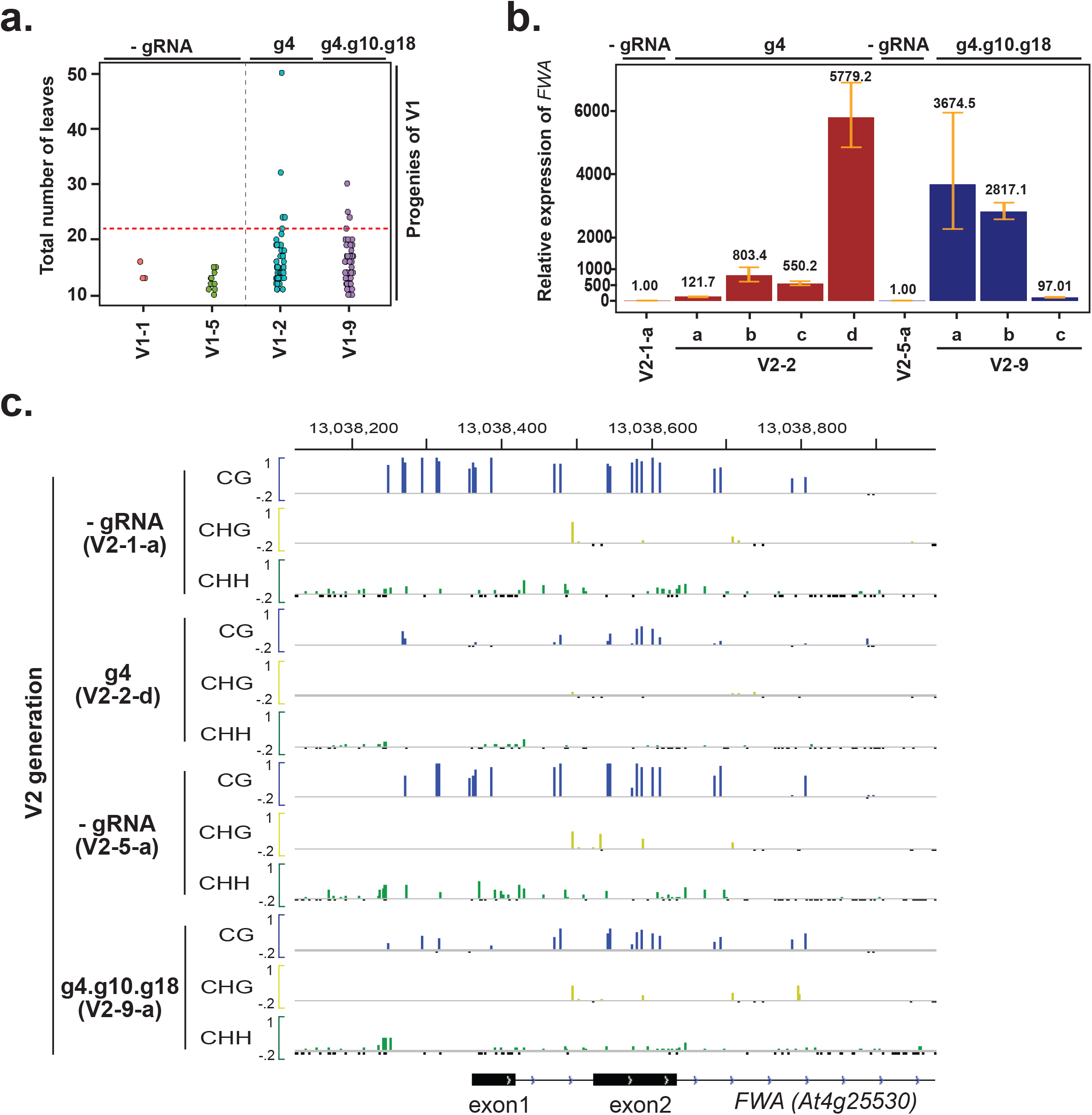
DNA demethylation and *FWA* expression is further enhanced in progeny of the TRV inoculated plants. (a) Dot plot showing flowering time in the progeny (V2) of virus inoculated plants (V1). Total numbers of leaves at the time of bolting are shown. (b) *FWA* expression in late flowering plants by reverse transcriptase-qPCR analysis. Error bars indicate standard deviations (n=3 technical replicates). (c) DNA methylation profile at the *FWA* promoter region in V2 plants descended from either g4 or g4.g10.g18-treated V1 plants (V1-2, V1-9), relative to matched progeny of -gRNA treated control (V1-1, V1-5) plants. Data is obtained from whole genome bisulfite sequencing. Colors indicate methylation context (CG=Blue, CHG=Yellow, CHH=Green). Only cytosines with at least 5 overlapping reads are shown. Small, negative values (black) indicate cytosines with 5 or more overlapping reads but no DNA methylation.

Next, we analyzed several V2 plants (schema in S4 Fig) for DNA methylation using the McrBC based DNA methylation assay at the *FWA* promoter as well as *FWA* expression by reverse transcriptase-qPCR. The late flowering V2 plants showed strong DNA demethylation at the *FWA* promoter relative to controls (S3b Fig), accompanied by a strong increase in *FWA* expression (Fig 3b). Effects on *FWA* expression were stronger in V2 than in V1, with *FWA* expression increased several thousand-fold in the three most demethylated V2 lines (V2-2-d, V2-9-a, V2-9-b). We also performed whole-genome bisulfite sequencing on the two V2 lines (V2-2-d, V2-9-a) with the strongest loss of DNA methylation (S3b Fig), as well as their respective controls (V2-1-a, V2-5-a). These data showed a strong reduction of CG methylation in the two V2 lines compared to controls (Fig 3c).

Since TRV can infect germline cells and be occasionally transmitted across generations [21, 29], we investigated whether TRV was present in the late flowering V2-plants. No PCR products specific to guide 4 sequences were detected in the V2 plants with delayed flowering (S3c Fig). These results suggest that *FWA* demethylation induced by TRV encoded guide RNAs is maintained in the subsequent generation in the absence of the guide RNAs.

### Near-complete loss of DNA methylation from *FWA* locus is attained in the progeny of the V2 plants inoculated with TRV encoded gRNAs

To analyze inheritance of *FWA* activation in the V3 generation, we selected three late-flowering V2-lines (V2-2-d, V2-9-a, V2-9-b) that showed a dramatic decrease in DNA methylation at *FWA* and strong upregulation of *FWA* expression, and analyzed flowering time in approximately 48 V3 progeny of each of the three lines. We also analyzed 12 plants from each of the two control -gRNA V2 lines (V2-1-a, V2-5-a). As expected, none of the progeny of control lines showed a late flowering phenotype (S1b Table). All three V2 lines gave rise to variable numbers of late flowering plants in the V3 generation, with the most efficient line producing 25% late flowering plants (Fig 4a and S1b Table). The observation that there was not 100% inheritance of late flowering in these lines is consistent with the dominant nature of *FWA* epigenetic alleles [12], as well as the incomplete demethylation of *FWA* in the V2 generation (Fig 3c).

**Fig 4.**
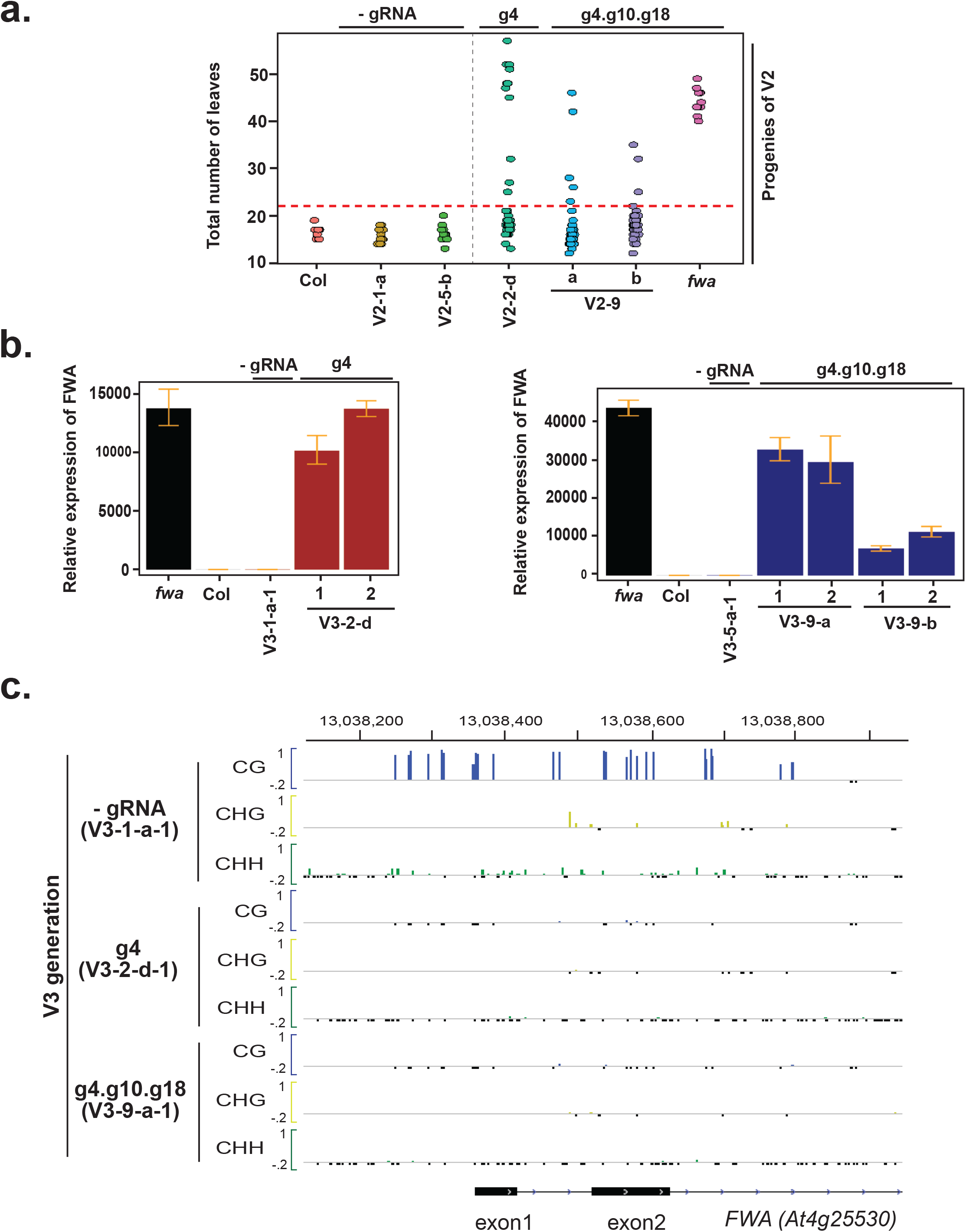
Near-complete loss of DNA methylation and strong gain of *FWA* expression in second-generation progeny of TRV-inoculated plants. (a) Dot plot of leaf count at flowering time of the second-generation progeny of TRV-inoculated plants (V3). (b) *FWA* expression in late-flowering plants by reverse transcriptase-qPCR. Error bars indicate standard deviations (n=3 technical replicates). Progeny of g4 inoculated plants shown left, g4.g11.g18 shown right. (c) DNA methylation profile at the *FWA* promoter region in V3 plants relative to control plants. Colors indicate methylation context (CG=Blue, CHG=Yellow, CHH=Green). Only cytosines with at least 5 overlapping reads are shown. Small, negative values (black) indicate cytosines with 5 or more overlapping reads but no DNA methylation.

To confirm that the late-flowering phenotype in the V3 plants is due to increased *FWA* expression, we performed *FWA* reverse transcriptase-qPCR (Fig 4b). *FWA* expression was increased several thousand-fold in all of the late-flowering lines, and in several plants, it was similar to the levels observed in an *fwa* epimutant (Fig 4b). We also performed whole-genome bisulfite sequencing in two different plants expressing *FWA* at high levels and found that these plants showed essentially complete DNA demethylation at the *FWA* locus (Fig 4c).

When TET1 was expressed in Arabidopsis at very high levels it was shown to cause genome-wide demethylation which indirectly led to demethylation and overexpression of *FWA* [27]. To rule out this possibility in our system, we analyzed genome-wide DNA methylation levels in V3 plants compared to respective controls. Similar to our previously published results [10], we observed little change in genome-wide DNA methylation levels in the V3 plants, suggesting that SunTag-TET1-mediated demethylation in these plants was restricted to the *FWA* locus (S5a and S5b Figs). Therefore, the TRV mediated *FWA* demethylation described here is highly specific with minimal off-target effects.

## Conclusion

In this study, we have developed a viral-based guide RNA delivery tool for CRISPR-mediated transcriptional activation and targeted DNA demethylation. We found that the targeted DNA demethylation can be heritable, even though viral RNAs are no longer detectable in progeny plants. Our TRV-tRNA-gRNA system gave 5 - 8% editing in the V2 generation in the best performing lines.

The tRNA-gRNA architecture used here has a few advantages over other guide RNA expression systems that have been used in viruses, such as the use of ribozymes or the use of multiple subgenomic RNA promoters [15, 18]. The presence of tRNA signal sequences was recently shown to facilitate heritable genome editing, which most likely also aided in the spread of TRV encoded gRNA sequences to increase the likelihood of activity in germ cells for epigenome editing. Additionally, in the tRNA-gRNA system, endogenous plant tRNA processing enzymes target the flanking tRNAs to produce mature guide RNAs allowing for the expression of multiple guide RNAs from the same virally-derived transcript [30]. The use of tRNA sequences in the viral RNA seems counter-intuitive as having these sequences would lead to the targeting of the viral RNA by the endogenous plant tRNA processing enzymes, which would theoretically reduce virus levels. However, the detection of viral RNA in the upper uninoculated leaves in this study (Fig 1b and 1d) and the detection of genomic edits in the progenies of TRV-tRNA-gRNA inoculated *Nicotiana benthamiana* plants in a recently published study [20], suggests that the TRV with tRNA sequences at least to some extent can evade the activity of plant tRNA processing enzymes in the inoculated cells.

Multiple plant generations were required to observe complete losses of methylation at the *FWA* locus when TRV encoded gRNAs were delivered to transgenic plants expressing the SunTag-TET1 system. In contrast, we previously observed that complete demethylation of *FWA* by SunTag-TET1 occurred in the first generation T1 plants when the construct encoded a ubiquitously expressed gRNA [10]. The most likely explanation for this difference is that the TRV-tRNA-gRNA system used here is less efficient at delivering gRNAs to all cells of the plant, resulting in chimerically demethylated plants. This is consistent with our analysis of DNA methylation of the first generation V1 plants in which we observed that only a fraction of the sequencing reads showed demethylation at CG sites. Partial demethylation in germ cells by TRV delivered gRNAs likely result in progressive multigenerational losses of methylation. This is consistent with several studies in plants that have shown that it can require multiple generations for stable and heritable establishment of methylation or demethylation [28, 31–33].

In summary, this study provides evidence for the utility of TRV to delivery gRNAs for CRISPR-CAS9 based epigenetic gene regulation.

## Materials and Methods

### Plant material growth conditions

The plants used in this study were grown under long-day conditions (16h light/8h dark) at 23°C in controlled growth chambers. SunTag-VP64 or SunTag-TET1 transgenic plants were selected on 1/2 MS medium + 35 μg/mL Hygromycin B (Invitrogen). The number of leaves at the time of flowering was calculated by counting the total number of rosette and cauline leaves.

### Plant inoculation

Rub inoculation of *Arabidopsis thaliana* plants (Columbia) was conducted using infected *Nicotiana benthamiana* leaves, 3 to 4 leaf staged *N. benthamiana* plants were agroinfiltrated with GV3101 strain containing the TRV RNA1 and RNA2(no guide RNA)/g4/g4.g10.g18 in a 1:1 proportion at an OD600 =1. Three to four days post inoculation, upper uninoculated systemically infected leaves were collected as inocula to infect Arabidopsis plants. Inoculated leaf tissue was used to rub-inoculate the two lowermost leaves of 12-14 day old Arabidopsis plants using 10 mM sodium phosphate. For agroinoculation of Arabidopsis plants, the protocol from Burch-Smith et al. was followed as detailed in the article [25]. Briefly, GV3101 strains containing either TRV RNA1 and RNA2(no guide RNA)/g4/g4.g10.g18 were mixed in 1:1 proportion at an OD 600 =1.5 and was agro-inoculated onto the lowest two leaves of 12-14 days old Arabidopsis plants.

### Cloning

#### Cloning sgPeBV promoter

Infectious cDNA clones of TRV genome (RNA1 and RNA2) pYL192 and pYL156 were obtained from Dr. David Baulcombe’s lab [34]. PeBV subgenomic coat protein RNA promoter was synthesized from SGI-DNA with XhoI and SmaI restriction sites at the 5’ and 3 ‘ ends of the promoter sequence, respectively. TRV RNA2 (pYL156) was digested with XhoI and SmaI. PeBV subgenomic promoter was cloned into vector backbones using the XhoI and SmaI sites by directional cloning.

#### Cloning of tRNA-guide RNAs into TRV with sgPeBV promoter

Plasmid pYL156 containing TRV RNA 2 sequence with sgPeBV promoter was digested with HpaI and SmaI. tRNA-guide4-tRNA and tRNA-guide4-tRNA-guide10-tRNA-guide18 sequences were synthesized by GenScript and were inserted at the HpaI and SmaI site by directional cloning.

### Quantitative real-Time PCR (qPCR) and reverse transcriptase-qPCR

Total RNA was extracted using Direct-zol RNA Miniprep kit (Zymo) from rosette and cauline leaves using in column DNase digestion. For reverse transcriptase-qPCR, cDNA synthesis was done using the SuperScript III First-Strand Synthesis SuperMix (Invitrogen). *FWA* transcripts were detected by using oligos JP7911 (5’-ttagatccaaaggagtatcaaag-3’) and JP7912 (5’-ctttggtaccagcggaga-3’). The ct values were first normalized to the housekeeping gene *ISOPENTENYL PYROPHOSPHATE:DIMETHYLALLYL PYROPHOSPHATE ISOMERASE 2* (*IPP2*) using oligos JP11859:(5′-gtatgagttgcttctccagcaaag-3′) and JP11860 (5′-gaggatggctgcaacaagtgt-3′). The delta ct Values were calculated relative to the control plants. Since *FWA* is not expressed in Columbia leaves for some samples, we got no cT values. To these samples we assigned a ct value of 40, which was equal to the maximum number of cycles run for the qPCR.

### McrBC–qPCR

Genomic DNA was extracted using CTAB method. For McrBC, 500 ng to 1 μg amounts of DNA were digested using the McrBC restriction enzyme for 4 h at 37°C, followed by 20 mins at 65°C. Equal amounts of DNA were used as controls by incubating them in buffer without the enzyme for 4 h at 37°C. qPCR of the *FWA* promoter was done using the oligos JP15049 (5′-*ttgggtttagtgtttacttg*-3′) and JP15050 (5′-*gaatgttgaatgggataaggta*-3′). The ratio between the digested to the undigested was calculated and was expressed relative to the control lines.

### Whole genome bisulfite sequencing analysis (WGBS)

CTAB-based method was used to extract DNA. Approximately 75 to 150 ng of DNA was used to synthesize whole genome bisulfite sequencing libraries using the Ovation Ultralow Methyl-seq kit (NuGEN). Raw sequencing reads were aligned to the TAIR10 genome using Bismark (48). Methylation ratios are calculated by #C/(#C+#T) for all CG, CHG, and CHH sites by a custom bash script. This was used to generate bigwig files for tracks to view on Integrative Genomics Viewer (IGV). Cytosines with more than 5 read coverage and zero methylation were represented by a negative black bar, while cytosines with less than 5 read coverage are represented by no bars. Reads with three consecutive methylated CHH sites were discarded since they are likely to be unconverted reads as described before (49). Whole genome metaplots were generated using the program ViewBS and following instructions of authors [35]. The same program was also used to calculate the genome wide methylation.

### Bisulfite PCR sequencing

CTAB-based method was used to extract genomic DNA and Qiagen EpiTect bisulfite kit was used to treat DNA per manufacturers instruction. Libraries were made from purified PCR products amplified by using previously described primers [6] using an Ovation Ultralow V2 kit (NuGEN). Libraries were sequenced on Illumina HiSeq 2500. Bisulfite PCR reads were mapped and processed using Bismark [36]. Unmethylated reads in CG and CHH contexts were obtained from Bismark output using a custom script (authors will provide upon request).

## Acknowledgements

We thank Dr. David Baulcombe for the gift of the pYL192 and pYL156 vectors, and Stephanie Zhou for technical help. We also thank Dr. Jason Gardiner and Dr. Ashot Papikian for the gift of SunTag-no guide VP64 and SunTag-no guide TET1 lines. We thank Mahnaz Akhavan for her help in Illumina sequencing. High-throughput sequencing was performed in the University of California, Los Angeles (UCLA) Broad Stem Cell Research Center BioSequencing Core Facility.

## Supporting information captions

**S1 Fig.**
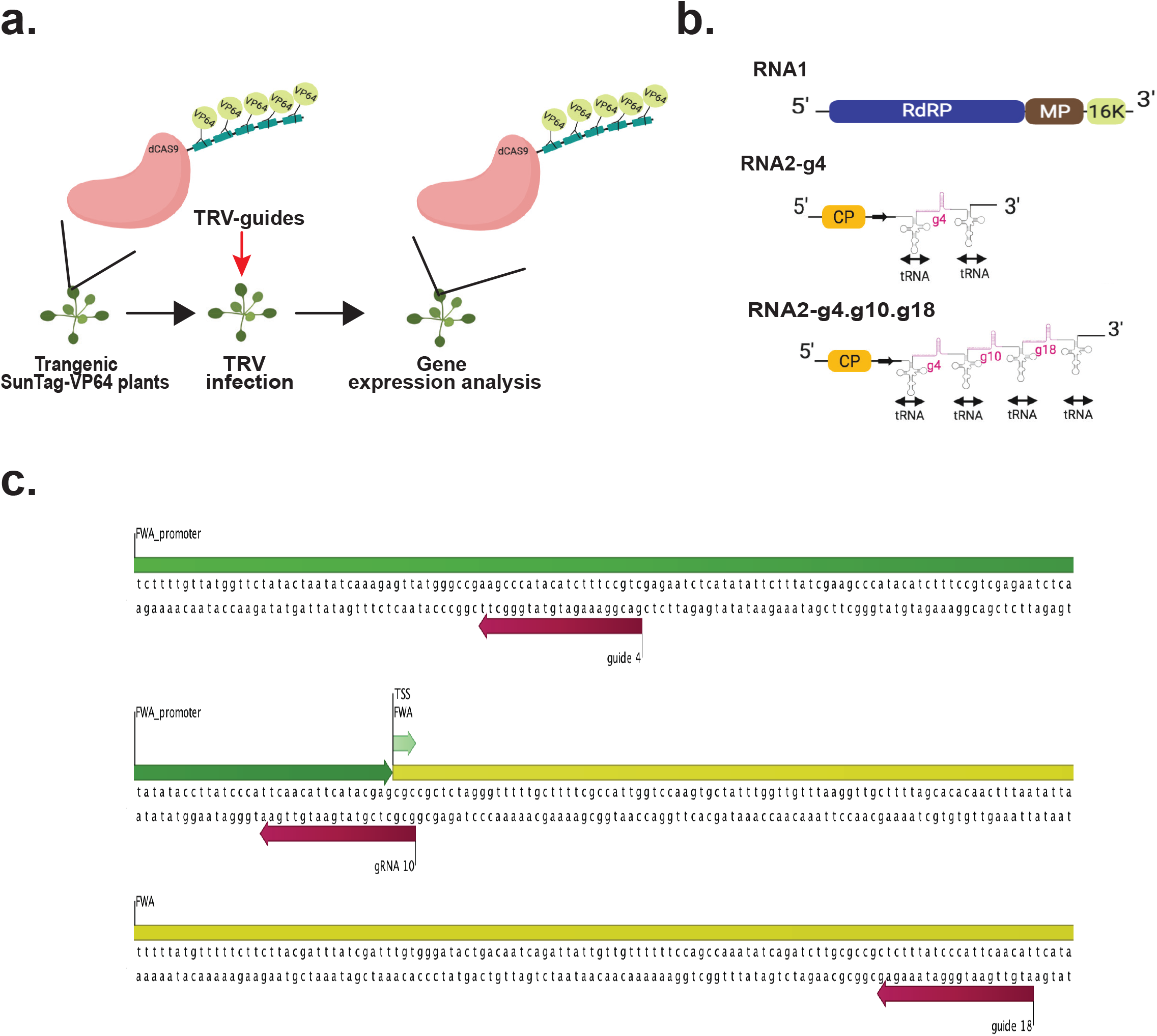
Details about the SunTag system and TRV-delivery guide RNA system used in this study. The *FWA* gene was targeted using guide RNAs delivered to a SunTag-VP64 overexpressing *Arabidopsis thaliana* line. (a) Schematic representation of the guide RNA delivery process by TRV to *Arabidopsis thaliana* plants. In the diagram, VP64 is shown as the effector, which is bound to the epitope tail. For targeted DNA demethylation experiments, plants with the SunTag system but with TET1 as the effector was used. (b) Sketch of TRV RNA1 and modified TRV RNA2 with a single guide RNA (guide 4(g4)) and three guide RNAs (guide 4, guide 10 and guide 18 (g4.g10.g18)) engineered into the genome. TRV RNA1 encodes viral RNA dependent RNA polymerase (RdRP), movement protein (MP), and suppressor of RNA silencing (16K). Guide RNAs were flanked by tRNA sequences to properly process the guide RNAs from the TRV RNA2 genome. Black arrow next to Coat protein (CP) denotes the sgPeBV promoter. (c) Location of guide RNA 4, guide RNA 10, and guide RNA 18 binding sites at *FWA* relative to transcription start site (TSS).

**S2 Fig.**
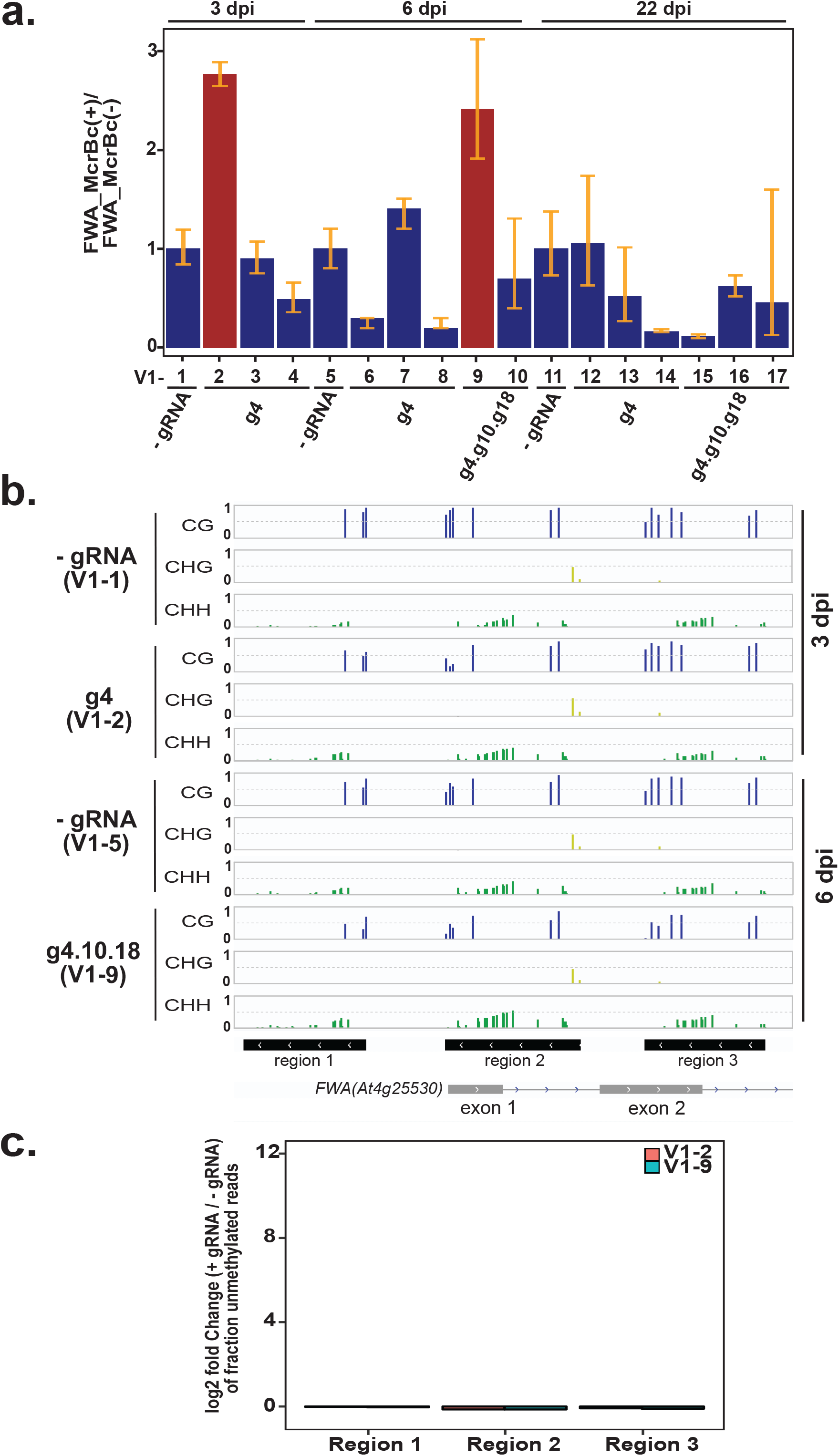
Selection of DNA demethylated lines at *FWA* promoter and the specific removal of CG methylation. (a) DNA methylation at *FWA* was analyzed by an McrBC-qPCR based assay for all plants in the V1 generation. Error bars indicate standard deviations (n=3 technical replicates). (b) DNA methylation data from bisulfite PCR sequencing, for cytosines in CG, CHG, and CHH contexts, at region 1 (Chr4:13038143 to 13038272), region 2 (Chr413038356 to 13038499), and region 3 (Chr4:13038568 to 13038695) of the *FWA* promoter. An average of 30,000 to 50,000 reads were obtained for each of the three regions. Colors indicate methylation context (CG=Blue, CHG=Yellow, CHH=Green). (c) Data in (b) is displayed as a bar graph of the log_2_ ratio of the fraction of fully CHH-demethylated reads in +gRNA plants relative to their no guide RNA controls.

**S3 Fig.**
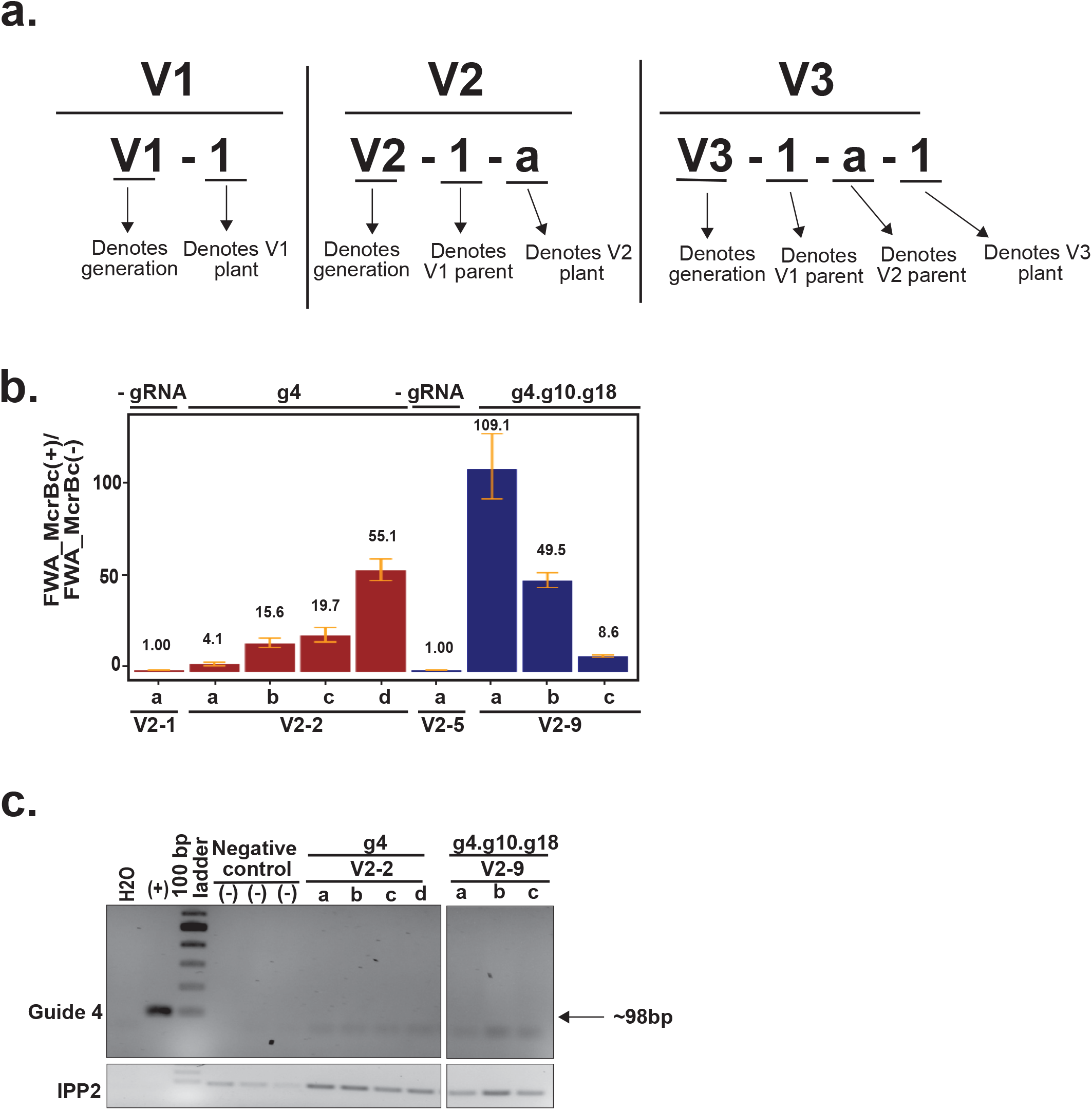
V2 generation plants exhibiting late flowering phenotype also display loss of DNA methylation. (a) Pattern of naming plants in V1, V2, and V3 generations. (b) DNA methylation at *FWA* by McrBC-qPCR based assay of plants exhibiting late flowering phenotypes. Error bars indicate standard deviations (n=3 technical replicates). (c) Agarose gel electrophoresis of reverse transcriptase PCR products amplified to detect guide RNA specific sequences in leaf samples of V2 plants exhibiting late flowering phenotype and their respective controls. H_2_0 = no-DNA negative control, (+) denotes positive control (plasmid containing guide 4). Negative controls are RNA from plants that were not inoculated with viruses but were grown side by side at the same time. IPP2 is used as quality control for the DNA sample.

**S4 Fig.**
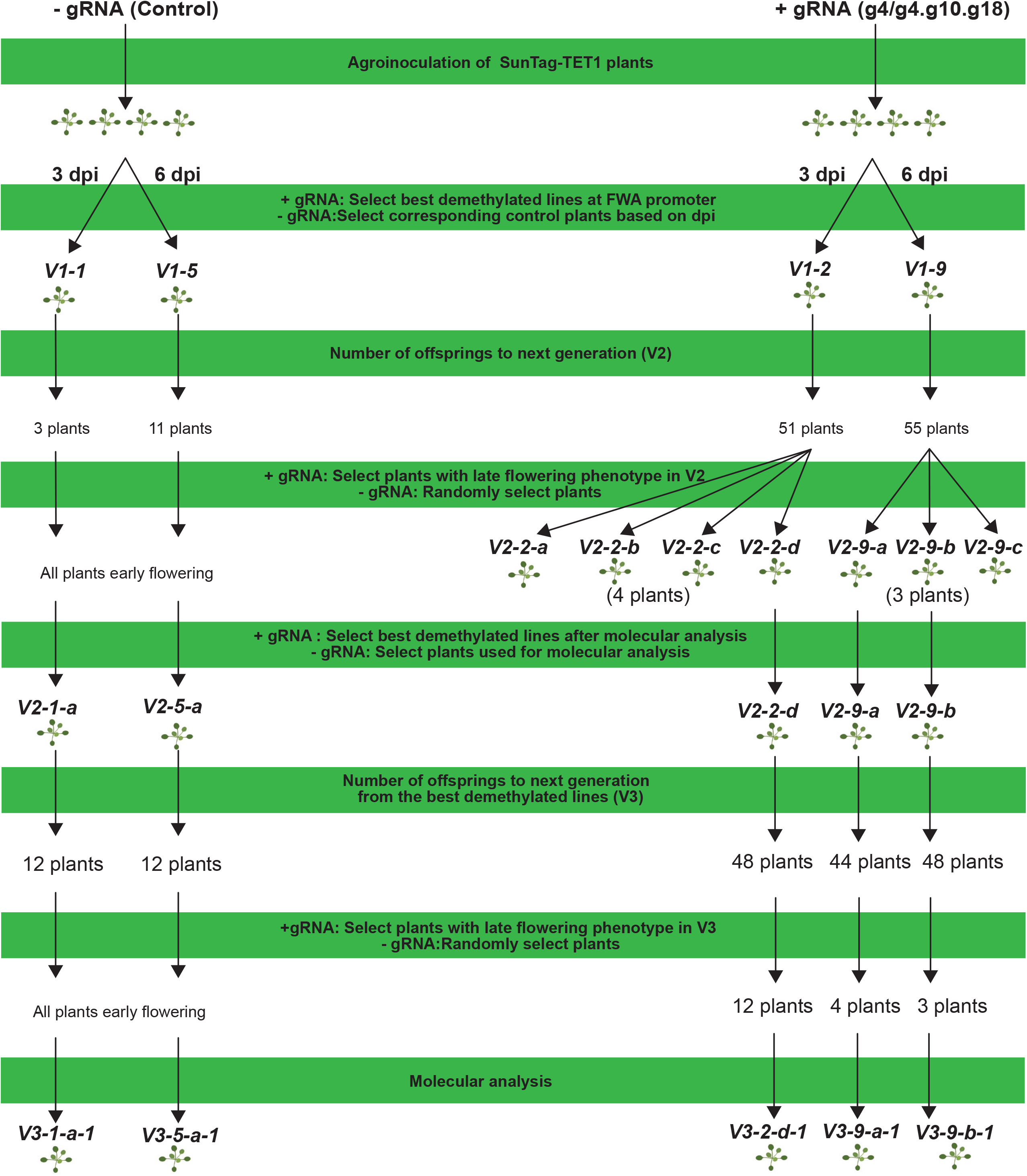
A schematic diagram depicting plant pedigrees and naming after viral delivery of guide RNA for targeted DNA demethylation.

**S5 Fig.**
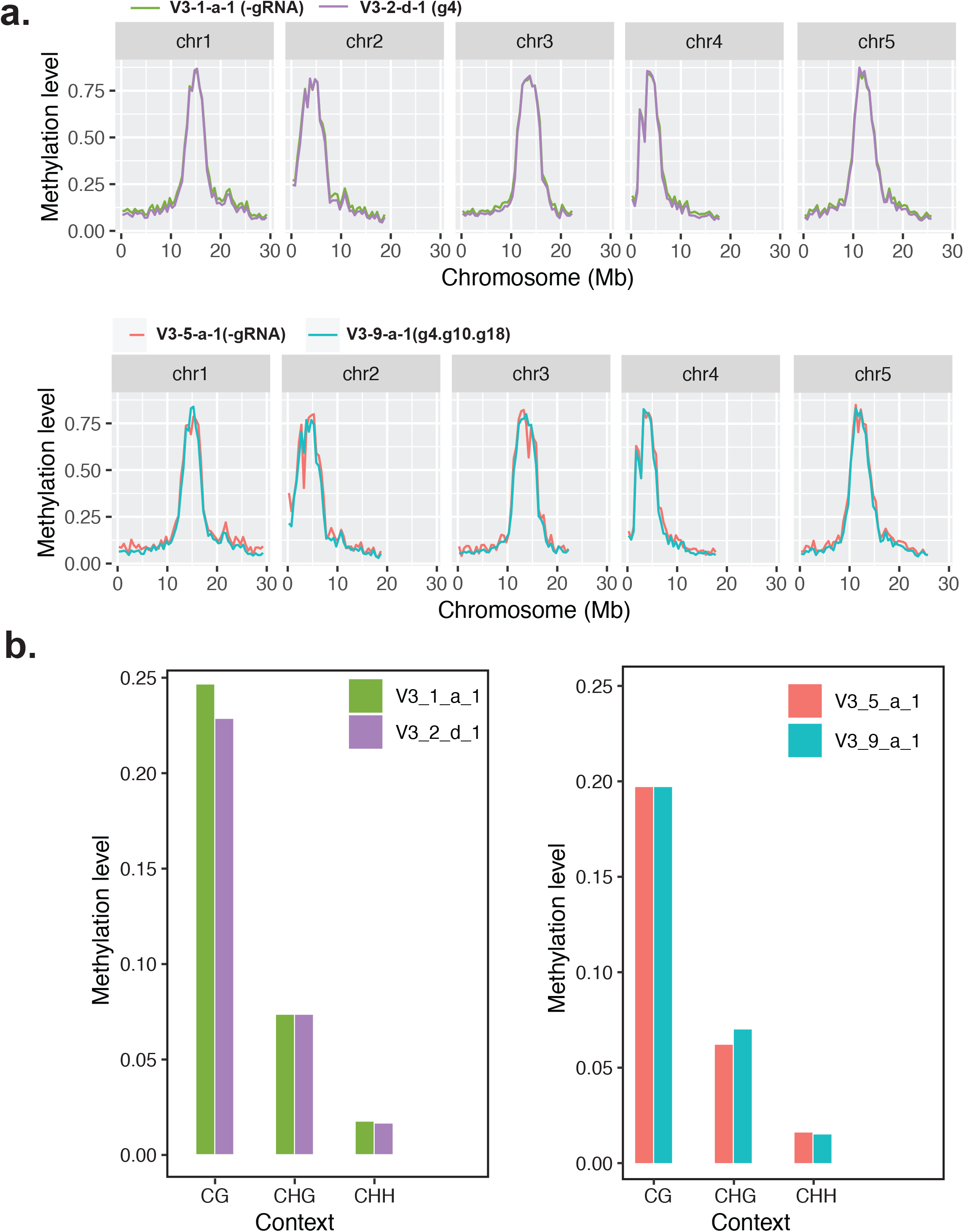
Late flowering plants in V3 generation exhibit minimal off target effects. (a) Whole genome CG methylation profile at the chromosome level in V3 plants exhibiting late flowering phenotype compared to their control plants. Metaplots over chromosomes were generated by ViewBS using default parameters. The lines selected for the analysis are shown at the top of the respective plots. (b) Bar graphs showing whole genome methylation percentage in two late flowering plants (V3-2-d-1, V3-9-a-1) compared to their matched controls (V3-1-a-1, V3-5-a-1).

**S1 Table.**
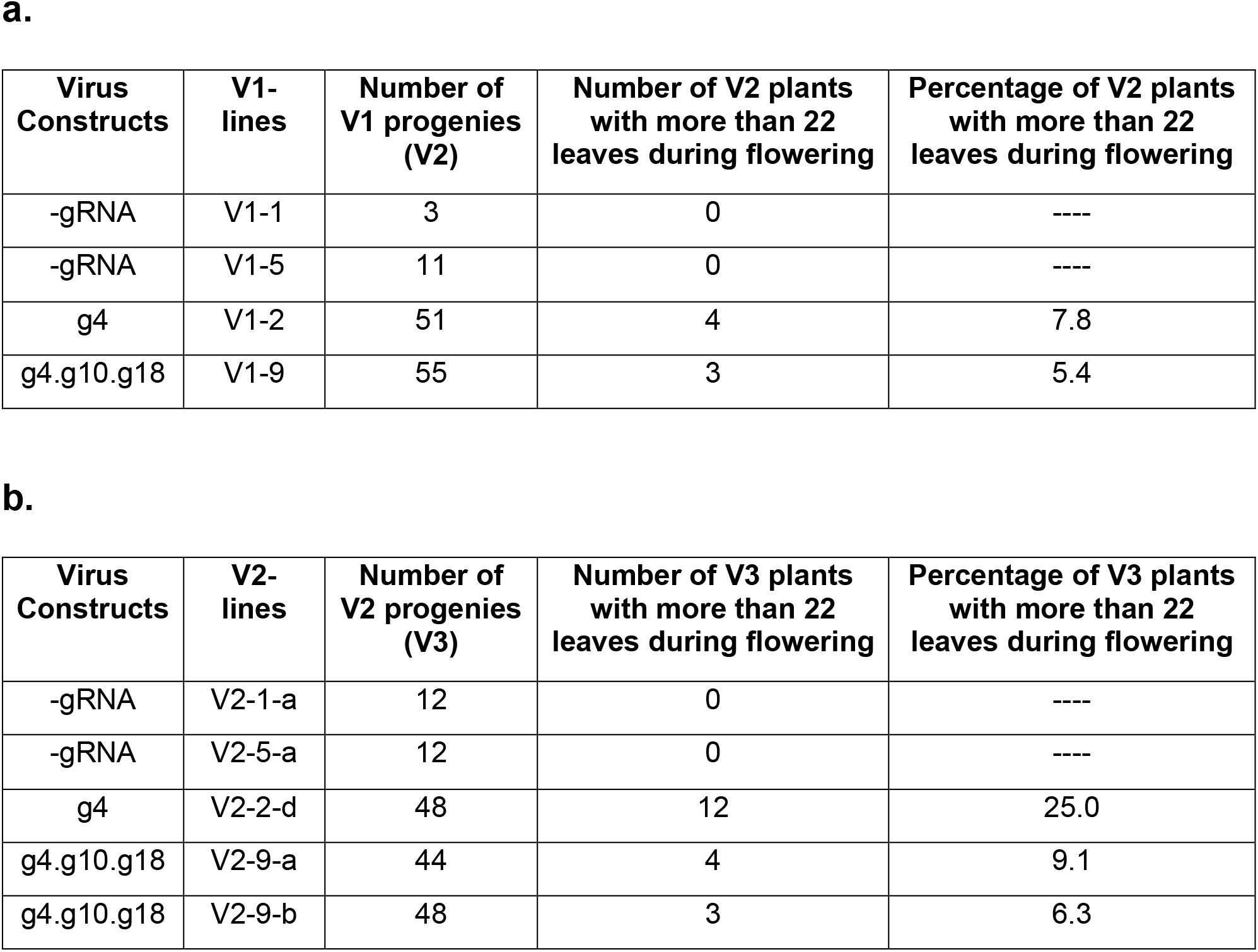
Details of Number of plants showing late flowering phenotype in the V2 and V3 generation. (a) Number of plants showing a late flowering phenotype in V2 generation. A late flowering phenotype was determined by counting rosette and cauline leaves at the time of bolting. Plants with more than 22 leaves at bolting were considered late flowering. (b) Number of plants showing a late flowering phenotype in V3 generation.

